# Skin Lesion Classification Via Combining Deep Learning Features and Clinical Criteria Representations

**DOI:** 10.1101/382010

**Authors:** Xiaoxiao Li, Junyan Wu, Hongda Jiang, Eric Z. Chen, Xu Dong, Ruichen Rong

**Author notes:** These two authors contributed equally (Xiaoxiao Li, Junyan Wu).

## Abstract

Skin lesion is a severe disease globally. Early detection of melanoma in dermoscopy images significantly increases the survival rate. However, the accurate recognition of skin lesion is extremely challenging manually visualization. Hence, reliable automatic classification of skin lesions is meaningful to improve pathologists’ accuracy and efficiency. In this paper, we proposed a two-stage method to combine deep learning features and clinical criteria representations to address skin lesion automated diagnosis task. Stage 1 - feature encoding: Modified deep convolutional neural networks (CNNs, in this paper, we used Dense201 and Res50) were fine-tuned to extract rich image global features. To avoid hair noisy, we developed a lesion segmentation U-Net to mask out the decisive regions and used the masked image as CNNs inputs. In addition, color features, texture features and morphological features were exacted based on clinical criteria; Stage 2 - features fusion: LightGBM was used to select the salient features and model parameters, predicting diagnosis confidence for each category. The proposed deep learning frameworks were evaluated on the ISIC 2018 dataset. Experimental results show the promising accuracies of our frameworks were achieved.

## 1 Introduction

Skin cancer is a major public health problem, with over 5,000,000 newly diagnosed cases in the United States every year. Melanoma is the deadliest form of skin cancer, responsible for an overwhelming majority of skin cancer deaths, and accounts for 75% of deaths associated with skin cancer [1]. In 2015, the global incidence of melanoma was estimated to be over 350,000 cases, with almost 60,000 deaths. Fortunately, if detected early, melanoma survival exceeds 95% [2]. For skin lesion imaging, dermoscopy is a noninvasive skin imaging technique of acquiring a magnified and illuminated image of a region of skin for increased clarity of the spots on the skin [3]. Dermoscopy technique was developed to improve the diagnostic performance of melanoma. The manual inspection from dermoscopy images made by dermatologists is usually time-consuming, error-prone and subjective (even well-trained dermatologists may produce widely varying diagnostic results)[3]. Nevertheless, the automatic recognition of melanoma from dermoscopy images is still a difficult task, as it has several challenges. First, the low contrast between skin lesions and normal skin region makes it difficult to segment accurate lesion areas. Second, the melanoma and non-melanoma lesions may have a high degree of visual similarity, resulting in the difficulty for distinguishing melanoma lesion from non-melanoma. Third, the variation of skin conditions, e.g., skin color, natural hairs or veins, among patients produce the different appearance of melanoma, in terms of color and texture, etc. Some investigations attempted to apply low-level hand-crafted features to distinguish melanomas from non-melanoma skin lesions [4]. Some researchers employed CNNs for melanoma classification, aiming at taking advantage of their discrimination capability to achieve performance gains [5, 6]. There is still much room to improve the accuracy of melanoma recognition by combining CNNs and clinical criteria. In this paper, we proposed a novel method based on deep convolutional neural networks and low-level image feature descriptors, which imitate clinical criteria representations, to solve skin lesion analysis towards melanoma detection problem.

## 2 Methods

### 2.1 Convolutional Neural Network

Skin lesions have large inter classes variations. Hence the key distinguishable features are difficult to be fully captured by traditional computer vision approach. Convolutional Neural Networks have led to breakthroughs in natural images classification and object recognition. Since CNNs have hierarchical feature learning capability, we modified ad fine-tuned state-of-the-art image classification networks — ResNet50 [7] and Dense201 [8] to encode the image features. The fully connected(FC) layers of both networks were modified by removing the last FC layer and adding FC layer with 128 kernels and FC layer with 7 (number of classes) for the 7—class skin lesions classification task.

Segmentation is an essential step for many classification approaches. Especially for skin lesion detection, background often contains noisy hair or camera masks. Accurate segmentation can benefit recognizing the skin lesion classes. We proposed a modified UNet [9] (architecture is shown in Appendix section) to generate lesion masks. Since the clear background is also useful for classification, instead of just masking out the foreground object, we firstly found the soft bounding box (lager than the smallest bounding box in order to include background) of the lesion. Then we crop the bounding box and padded it to its smallest square with 0s, as shown in Fig. 1.

**Fig. 1.**
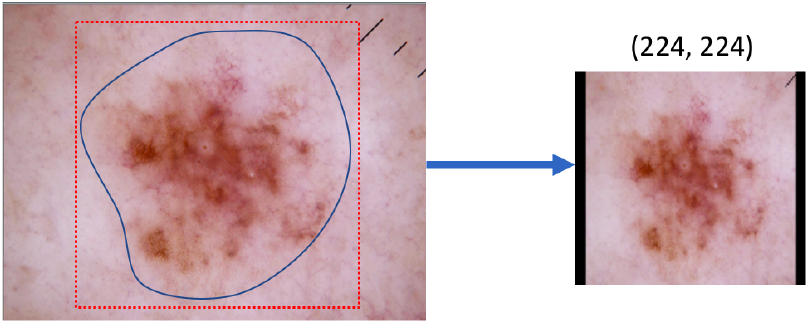
Finding bounding box, cropping and padding for the soft-segmented object

### 2.2 Clinical Criteria Representations

In this subsection, we explain the features generated with traditional computer vision methods. The feature descriptors tried to imitate clinical diagnosis procedure. Common skin diseases are primarily diagnosed by observation. In the diagnosis process, doctors make the diagnosis and provide suggestions for treatment according to the visual information and the empirical criteria. In practice, there exist clinical criteria used for diagnosis, which cover the major characteristics of skin lesions and can help dermatologists to make an accurate diagnosis, including color features, texture features, and morphological features. For the color feature, we generated the RGB and HSL features and what proportion of the image was that color. For the texture features, we produced SIFT and LBP features. For the morphological features, we calculated *solidity* and *circularity*, *image ration* and *area ratio.* Firstly, we detected a convex hull for the object region. *Solidity* was defined as the ratio of pixels in the region to pixels of the convex hull images. Then we approximated the mask contour as a line through the centers of border pixels using a 4-connectivity, notated as *perimeter.* We defined *circularity* as the ratio of the region area to *perimeter*^2^. Also, we found the rectangle bounding box for the region. *Image ration* was defined as the ratio of bounding box area to image area and *area ratio* was defined as the ratio of region pixel area to the bounding box area.

### 2.3 Feature Fusion

We used LightGBM[10] to combine the traditional features and CNN features. The LightGBM is a boosting tree-based learning algorithm. It provides multiple hyper-parameters for achieving best performance. It outperforms most state-of-art machine learning algorithms.

## 3 Experiments and Results

### 3.1 Dataset

The dataset we used in this challenge consisting of 10015 images (327 actinic keratosis, 514 basal cell carcinoma, 115 dermatofibroma, 1113 melanoma, 6705 nevus, 1099 pigmented benign keratosis, 142 vascular lesions) extracted from the ISIC 2018: Skin Lesion Analysis Towards Melanoma Detection grand challenge datasets [12, 13]. Each data is RGB color image, with size 450 × 600.

From the data with ground truth, We split 70% data as training set, 10% as the validation set, which was used to find the early stopping epoch and 20% as testing set to evaluate our algorithms.

As the number of images in each category varies widely and seriously unbalanced, we accordingly augmented the images of different classes in the training set. The augmentation methods included randomly left/right rotation (maximum 25°), left-right flipping, top-bottom flipping and zoom-in cropping with ratio 0.8. All the input image was resized to (224,224) in our application.

### 3.2 Algorithm Pipeline Implementation

The flowchart of our pipeline is shown in Fig. 3. In stage 1, we generated image features through deep learning and traditional handcrafted approaches. We used CNNs to encode the images features, meanwhile capture experimental features by descriptors. In the CNN feature encoder, we used ResNet50 and DenseNet201 to encode both raw RGB images and image patches after crop-padding. The loss function we used to train the networks is categorical cross-entropy. Since DenseNet201 could not handle the raw images inputs well, based on our testing experiment, we excluded this feature from the CNNs feature encoder. Hence, the CNNs feature encoder generated 3 groups of features: 1) raw images encoding with ResNet50; 2) masked images encoding with ResNet50 and 3) masked images encoding with DenseNet201. Each feature was represented as on 1D 128–dim vector, which is the output of FC(128) of the CNNs. From the color feature descriptor, we generated RGB and HSL features, which both are 3×1 vector. Also, we calculated the proportion of R channel, which is a scaler in [0, 1]. From the texture descriptor, we generated the SIFT features as 384×1 vectors and LBP feature as 27×1 vectors. From the morphological feature descriptor, we generated *solidity* and *circularity, image ration* and *area ratio,* which are 4 scalers.

**Fig. 2.**
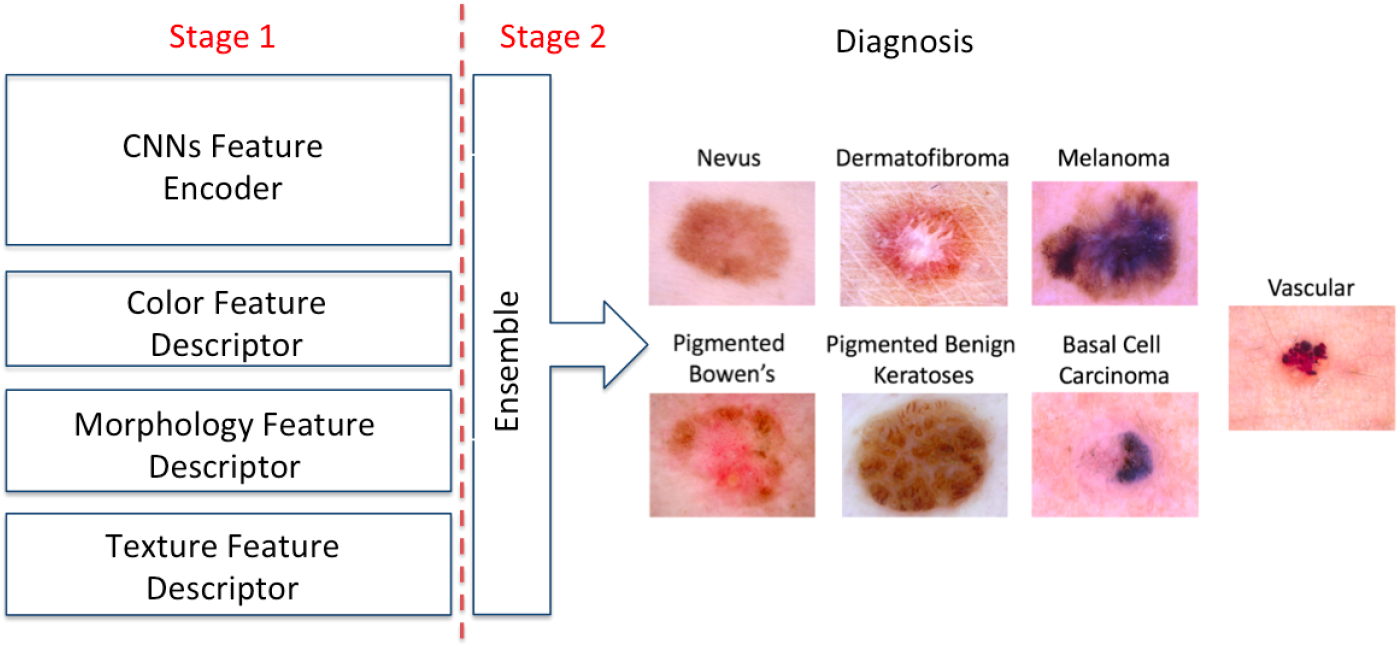
The flowchart of the proposed framework

In stage 2, we rescaled the features from stage 1. Then we used LightGBM combining the features and selected the best ensemble parameters. To train the LightGBM, we use one-vs-all (OvA) loss function [11].

### 3.3 Evaluation

#### Experiment Results on Training Dataset

the stopping epoch then tested our ensemble model on the testing set. The performance was summarized in Table 1.

**Table 1.** Summary of performance in the training data

| Categories | Precision | Recall | F1 score | Samples |
| --- | --- | --- | --- | --- |
| MEL | 0.62 | 0.79 | 0.66 | 223 |
| NV | 0.95 | 0.91 | 0.93 | 1341 |
| BCC | 0.75 | 0.85 | 0.80 | 103 |
| AKIEC | 0.63 | 0.68 | 0.66 | 66 |
| BKL | 0.73 | 0.76 | 0.75 | 220 |
| DF | 0.78 | 0.78 | 0.78 | 23 |
| VASC | 0.83 | 0.83 | 0.83 | 29 |
| Total | 0.86 | 0.85 | 0.86 | 2005 |

#### Compare Different Model on Training and Validation Dataset

We compared different input schemes and model as listed in Table 2 on the training and validation dataset. For the training dataset, it was evaluated on the same split testing set as mentioned above. The results of the validation dataset were achieved by submitting prediction results to the scoring system. The decimals in the table are the normalized multi-class accuracy for each case.

**Table 2.** Comparing different model strategies

| Model | ResNet | ResNet<br>+Crop | DenseNet<br>+Crop | Clinical<br>Feature | **All-**<br>**Fusion** |
| --- | --- | --- | --- | --- | --- |
| Training | 0.822 | 0.798 | 0.845 | 0.472 | **0.855** |
| Validation | 0.836 | 0.848 | 0.762 | 0.319 | **0.853** |

We have observed that the fusion model achieved better and more stable performance on normalized multi-class accuracy (the ranking metric), which demonstrates the effectiveness of the LightGBM fusion scheme.

## 4 Conclusion

In this paper, we proposed an ‘CNNs features + clinical criteria representations’ ensemble method for skin lesion analysis towards melanoma detection. Our combination approach and training strategies enlarged the ability of automatic lesion classification model to capture the reliable features. The LightGBM feature fusion method increased the robustness by randomly searching the parameters and selecting the salient features. Extensive experiments conducted on the open challenge dataset of Skin Lesion Analysis Towards Melanoma Detection (ISIC 2018) demonstrated the effectiveness of the proposed method. Our method combining deep CNNs and traditional computer vision features based on clinical criteria with effective training mechanisms can be employed to solve complicated medical image analysis problems, even with limited and unbalanced training data. Further work will include investigating integrating probability map generated by segmentation in our networks and explore more applications.

## Appendix

**Fig. 3.**
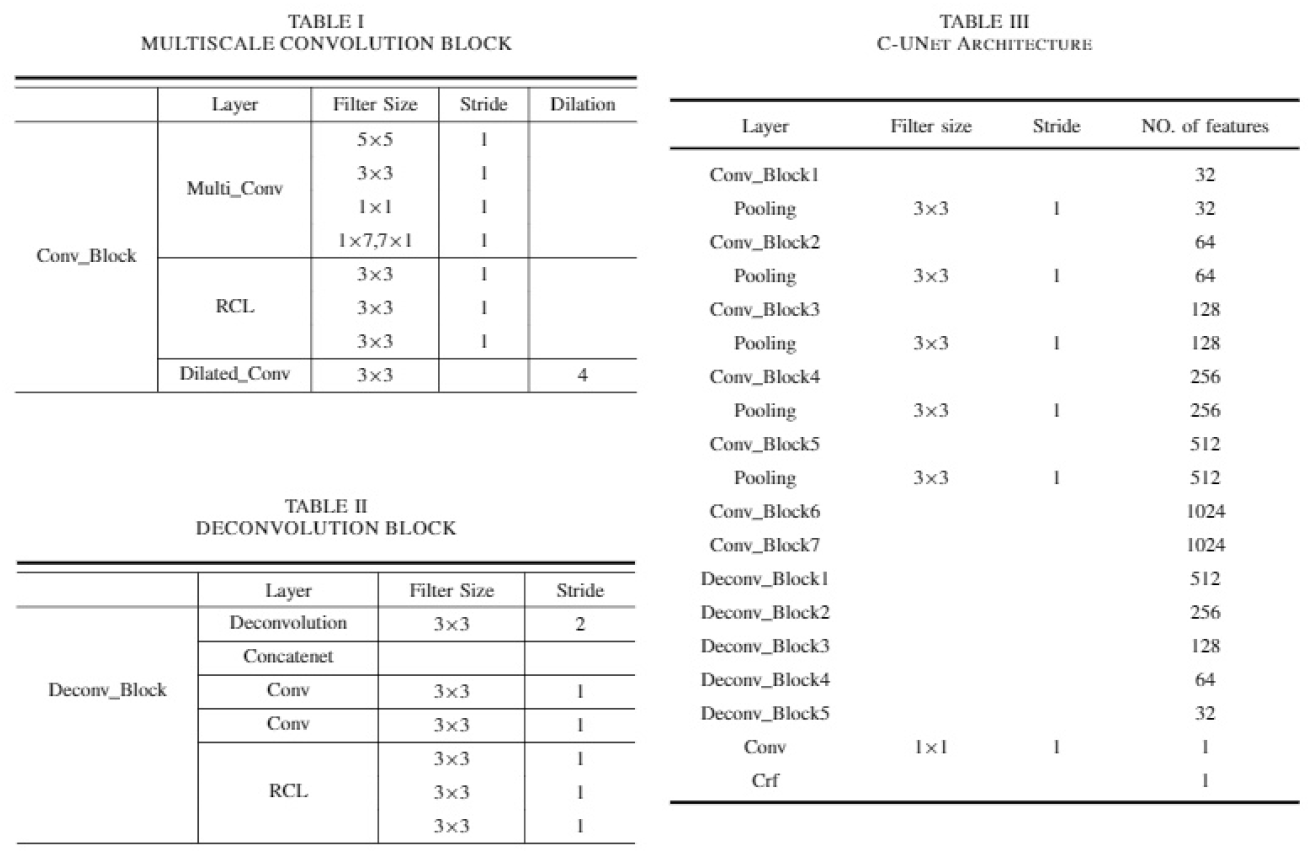
Segmentation Network

